# Improved cloning-free one-step CRISPR-Cas12a-assisted tagging of mammalian genes using PCR generated reagents (PCR tagging)

**DOI:** 10.64898/2025.12.02.690677

**Authors:** Dan Lou, Daniel Kirrmaier, Melanie Krause, Shengdi Li, Krisztina Gubicza, Konrad Herbst, Matthias Meurer, V. Talya Yerlici, Omar M. Wagih, Lars M. Steinmetz, Michael Knop

**Affiliations:** Zentrum für Molekulare Biologie der Universität Heidelberg (ZMBH), Im Neuenheimer Feld 345, 69120 Heidelberg, Germany; Deutsches Krebsforschungszentrum (DKFZ), DKFZ-ZMBH Alliance, Im Neuenheimer Feld 280, 69120 Heidelberg, Germany; EMBL, Genome Biology Unit, Meyerhofstr. 1, 69117 Heidelberg, Germany; SyntGen Lab, Im Neuenheimer Feld 345, 69120 Heidelberg, Germany; Orb Therapeutics, 1300-661 University Ave. Toronto, Ontario Canada M5G 1M1

## Abstract

Precise DNA integration in mammalian genomes using CRISPR/Cas based strategies remains challenging due to competing DNA repair pathways. We present an update for our CRISPR/Cas12a-assisted PCR-tagging method using linear donor cassettes and self-targeting gRNAs for gene tagging. By incorporating 2A-linked selection markers and chemically modifying PCR cassettes with phosphorothioate and biotin, along with the inhibition of NHEJ, we improve precise insertions while reducing unwanted integration products and we devise a variant of the method for seamless gene tagging. To profile integration outcomes, we further developed our integration site sequencing protocol (Tn5-Anchor-Seq, Meurer et al., 2018) to obtain a protocol using long-read (ONT) sequencing coupled with a custom computational pipeline. This enables high-resolution quantification of all integration events with respect to integration by HDR or NHEJ, as well as the presence of concatemers and off-target integrations. Our data reveal gene-specific editing profiles and demonstrate that combining cassette design with DNA repair modulation yields up to 5-fold increases in HDR and suppression of concatemers and indels. Together, our improvements outline an robust strategy for efficient and high-fidelity gene tagging in mammalian cells, facilitating functional genomics and cell engineering applications.

## Introduction

The introduction of CRISPR/Cas systems in genetic engineering has transformed both fundamental biological science and therapeutic research (Anzalone et al., 2020; Doudna, 2020). Among many different applications, the CRISPR endonucleases Cas9 and Cas12a have facilitated the site-specific insertion of foreign DNA into genomes. They allow programmed double-strand breaks (DSBs) in DNA, utilizing short RNA transcripts to direct the nucleases. Heterologous DNA repair templates possessing free ends can be incorporated into the genome by cellular DNA repair pathways, often via non-homologous end joining (NHEJ), which may utilize short microhomologies at the DNA ends. Alternatively, integration can also use homology directed repair (HDR) if the repair template provides longer stretches of sequence homology between the repair template and the selected genomic locus. When attempting to integrate a piece of DNA precisely at a specific position in the genome, alternative DNA repair pathways and variability in processing the DNA ends before integration compete with each other, leading to a variety of different outcomes (reviewed in: Averina et al., 2024; Xue & Greene, 2021). This can substantially hamper the success of targeted insertion (e.g. the construction of gene-fusions with fluorescent protein reporters) because the probability of perfect insertion at both ends of the reporter are very low (Koch et al., 2018). Further complications arise from concatemerization of the linear repair template (Folger et al., 1982, 1985); these constitute unwanted side-products that further lower the efficiency of the desired reaction. Nevertheless, specific modifications of the procedures along with tailored strategies to generate the repair template in a manner that favours the desired outcomes, e.g. using chemically modified DNA (Gutierrez-Triana et al., 2018), has led to a plethora of methods to introduce foreign DNA into a genome. These approaches, for example, enable the creation of gene-reporter fusions for various downstream functional studies (such as GFP tagging for protein dynamics). These methods usually combine genetic constructs (e.g. plasmids, viral vectors) harboring the donor DNA along with genetic constructs for the expression of the gRNAs and/or the Cas-nuclease proteins. Alternatively, some protocols include direct transfection of purified components, e.g. gRNAs or even premade cgRNA/Cas protein complexes (Anguela & High, 2019; Choi et al., 2023; Hale et al., 2009; Lackner et al., 2015; Saito & Kanemaki, 2021; Schmid-Burgk et al., 2016; Philip et al., 2025). The availability of such methods for gene tagging has greatly facilitated functional gene studies (Huh et al., 2003; Kanca et al., 2017; Serebrenik et al., 2024), creation of disease models (H. S. Kim et al., 2024; Shalem et al., 2015), and clinical therapeutic applications (Cetin et al., 2024; Li et al., 2020). Most methods for gene tagging involve gene-specific cloning steps and the production of relatively costly resources (protein, RNA), and thus studies where systematic tagging of larger numbers of genes is conducted, are rare (Cho et al., 2022; Husser et al., 2024; Kim et al., 2024; Roberts et al., 2017).

In yeast, gene tagging is commonly performed using PCR-generated linear DNA fragments that contain flanking homologous sequences, the tag, a heterologous terminator and a selection marker. This strategy is highly efficient, with tagging-efficiencies approaching 100% and fidelities exceeding 80%, even without CRISPR/Cas components, due to the predominance of HDR pathways. This makes ‘PCR tagging’ easily scalable and enables the creation of genome-wide resources of tagged strains (Janke et al., 2004; Mathur & Kaiser, 2014; Miao et al., 2024). Upon inclusion of a CRISPR/Cas cut, the fidelity of tagging increases to >95% and the number of obtained clones increases up to 1000-fold enabling the construction of genome-wide one-step pooled gene-tagging clone libraries (Buchmuller et al., 2019).

In contrast, mammalian cells present a very different repair landscape. Foreign DNA is predominantly integrated through error-prone repair mechanisms such as classical NHEJ (c-NHEJ) and microhomology-mediated end joining (MMEJ) (Scully et al., 2019; Yeh et al., 2019), rather than by the more precise HDR pathway. Consequently, integration events often deviate from the intended HDR-mediated design, leading to unintended mutations, insertions or deletions, off-target integrations, and concatemeric insertions (Erbs et al., 2023; Riesenberg et al., 2023; Suchy et al., 2024; Zou et al., 2023).

Towards the goal of developing a universal gene-tagging method for mammalian cells, we adapted the PCR-tagging strategy from yeast and demonstrate that simple PCR-generated linear DNA fragments can be used for efficient gene tagging in mammalian systems (Fueller et al., 2020). This involves a PCR product that guides its own integration into the genome by expression of a gRNA for Cas12a cleavage and two homology arms (HA) for HDR. The two PCR primers provide all gene-specific elements, and include in addition a protospacer sequence for a gRNA targeting the insertion site. All functional elements such as the tag-sequence including a terminator or an additional selection marker are provided by a generic template (Fueller et al., 2020). With this ‘mammalian PCR tagging’ method constitutes a simple and rapid gene tagging strategy in mammalian cells that has been widely used. Nevertheless, it does not yet come anywhere close to the efficiency and fidelity of the yeast PCR tagging method, thus leaving room for improvements. In addition, a systematic understanding of the resulting integration outcomes is essential for refining knock-in experiments, improving editing efficiency to achieve predictable and precise outcomes, and to efficiently isolate clones with correct editing outcomes.

To this end, improved selection strategies, enhanced precision in genomic integration, and convenient ways to analyze tagging outcomes are needed. In our current approach, selection markers are placed on the PCR cassette downstream of the tag and a polyA signal. This strategy enriches cells carrying the marker, but it does not ensure that the marker and tag are correctly inserted at the destination locus. Furthermore, next-generation sequencing-based methods have been developed to analyze the outcomes of such gene-integration events by amplifying the insertion junctions of the foreign DNA (Buchmuller et al., 2019; Fueller et al., 2020; Schmid-Burgk et al., 2016). Bioinformatic tools such as CRISPResso2 (Clement et al., 2019), ampliCan (Labun et al., 2019), and Knock-knock (Canaj et al., 2019) are optimized for short-read sequencing data, where they detect mutations, indels, off-target events, or simple HDR/NHEJ outcomes. Long-read-based tools such as DAJIN (Kuno et al., 2022) have been developed, but their ability to resolve the full spectrum of complex integration patterns typical of gene tagging is limited. Therefore, a comprehensive tool that leverages long-read sequencing to systematically characterize diverse knock-in outcomes remains lacking.

In this work we address these gaps and employ selection markers fused in-frame to the tag of choice (e.g. mNeonGreen) using a 2A peptide spacing the two protein domains ensures that the selection marker is cleaved and set free to function in the cytoplasm (Schmid-Burgk et al., 2016). We furthermore explore the use of various chemical inhibitors and modified oligos to prevent processing the repair template by DNA damage repair pathways that produce unwanted outcomes. We also developed a versatile deep sequencing platform (Tn5-Anchor-Seq), coupled with a computational pipeline, to characterize and quantify various integration outcomes following our adopted CRISPR-Cas12a-assisted PCR tagging strategy. Together, our work describes robust PCR-based procedures for gene tagging using generic terminators or seamless tag insertion for conducting more streamlined and fast-paced protein localization experiments.

## Results

### Improved selection and characterization of integration outcomes of CRISPR-Cas12a-assisted PCR tagging

Our previous PCR tagging protocol suffers from inefficient selection yielding between 20% and 60% of cells exhibiting a fluorescent signal (in the case of GFP tagging) at the correct location (Fueller et al., 2020), leaving room for improvements. To improve selection of correctly tagged clones we inserted a self-cleaving 2A peptide sequence followed by the phosphotransferase II (nptII) gene that confers resistance to neomycin or G418 (geneticin) at the C-terminus of fluorescent proteins (Figure 1a). This in-frame marker strategy has been developed previously and found to improve selection of correctly tagged genes in other tagging methods (Schmid-Burgk et al., 2016). Using this cassette for PCR-tagging (Figure 1a) in HEK293T-N cells (HEK293T cells where the neomycin gene linked to the T-antigen has been deleted), we tagged HNRNPA1 with mNeonGreen and observed tagging efficiencies of 70% and more (Example in Figure 1b), compared to 32% when employing a cassette in which the selection marker was inserted as a separate gene (Fueller et al., 2020). These results highlight the benefit of this strategy.

**Figure 1.**
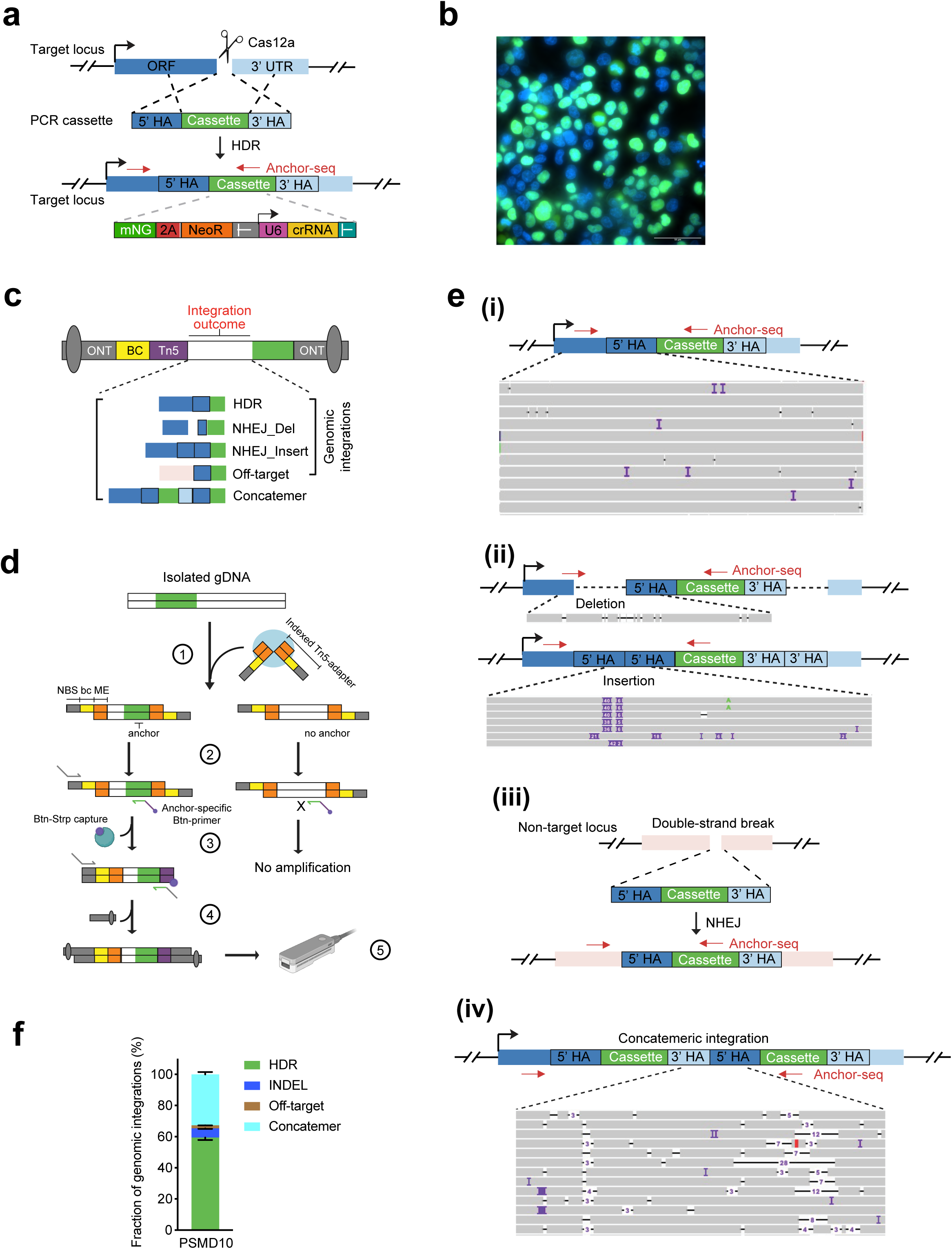
Mapping genomic integration outcomes following CRISPR/Cas12a-assisted PCR-tagging by Tn5-Anchor-Seq. (**a**) Schematic of PCR tagging strategy: The tagging cassette contains the tag-2A-marker fusion which is followed by a polyA adenylation site for proper transcript termination. The cassette also contains a functional gRNA gene. The protospacer sequence of the gRNA is provided by one of the PCR primers, thus allowing the use of generic PCR templates with gene-specific primers (Fueller et al., 2020). Upon transfection the gRNA gene is expressed and targets Cas12a (cotransfected on a plasmid) to induce a DNA DSB at the target location, followed by integration of the PCR cassette via homology-directed repair (HDR) or alternative pathways. (**b**) Representative microscopy image from HEK293T-N cells showing HNRNPA1 tagging. mNeonGreen fluorescence and HOECHST staining are shown. Images acquired using Nikon Ti-E widefield epifluorescence microscope with a 60x objective. Scale bar, 50 µm. (**c**) Schematic of sequencing read and integration outcomes. (**d**) Experimental workflow of Tn5-Anchor-Seq: 1) Isolated genomic DNA (gDNA) was fragmented using Tn5 transposase, which was loaded with indexed adapters comprised of a Tn5 mosaic end (ME), an 11-bp index (bc), and a nanopore sequencing primer binding site (NBS). 2) DNA fragments containing the anchor sequence were PCR-amplified with biotinylated primers specific to anchor sequence and adapter-specific primers directed at the NBS. 3) Biotinylated PCR products were enriched via a biotin-streptavidin capture step and nanopore barcodes and adapters were introduced by a nested PCR. 4) Final anchor-specific amplicons were sequenced on a Nanopore MinION platform. (**e**) Schematics illustrating genomic integration scenarios following PCR tagging: i. HDR-mediated integration; ii. NHEJ-mediated deletions and insertions; iii. Off-target integration; iv. Concatemeric integration. Corresponding IGV snapshots show representative sequencing reads for the indicated regions. (**f**) The frequencies of genomic integrations at *PSMD10* locus.

Despite the high fraction of cells exhibiting correctly localized signals, it is unclear what fraction of insertions occurred via HDR, and which via other pathways such as NHEJ that still may lead to in-frame fusion of the tag with the target protein. In addition, multiple PCR fragments in the form of concatemeric DNA might have been integrated (Figure 1c). To characterize the results of such gene-tagging experiments we next adopted our anchor-seq pipeline (Buchmuller et al., 2019; Fueller et al., 2020) for a full characterization of all possible outcomes of such tagging experiments. Compared to the previously published method, the optimized protocol replaces genomic DNA fragmentation using ultrasound with tagmentation using Tn5 (see Methods). The tagmentation adapter contains DNA barcodes to enable sample pooling after tagmentation. Biotinylated adapter-specific primers for PCR are used to enhance specificity through a biotin-streptavidin capture (Figure 1d). A second nested PCR using generic primers enables specific and sensitive amplification of the junctions encompassing neighbouring DNA flanking the 5’-homology arms of the PCR cassettes, followed by sequencing of the amplicons using Oxford Nanopore Technology (ONT) sequencing.

Inspection of the obtained reads revealed several different types of integration outcomes where the PCR cassette was inserted in different ways at the DSB site. We observed the expected homology-directed repair (HDR) outcomes where the donor-encoded tag (mNeonGreen) was scarlessly installed (Figure 1e(i)). NHEJ-mediated insertions where the mNeonGreen sequence was ligated directly to the genomic DNA yielded heterogeneous outcomes, depending on different outcomes of the DNA trimming during the repair process. Usually, these events were characterized by a duplication of parts of the homology arm (Figure 1e(ii)). All fusion events identified by sequencing must be in-frame so that the tag and associated selection marker are still expressed. However, the high error rate of ONT sequencing (of 3-5%, including small indels (1-10bp, often occurring in homopolymeric regions)) precludes a more detailed analysis of individual events. We also observed off-target outcomes where mNeonGreen was inserted into unintended locations in the genome (Figure 1e(iii)). These events are independent of Cas12a and occur occasionally when linear DNA is transfected into cells (Fueller et al., 2020) caused by random genomic integrations (Folger et al., 1982, 1985). We do not know whether these insertions do lead to the expression of the tag (which is necessary for selection) or whether these are hitchhike insertions in the background of cells containing a successful tag insertion. We also observed concatemeric knock-in outcomes in which multiple copies of the PCR cassette were ligated together (Figure 1e(iv)). We previously found that in some cases these concatemers can accidentally lead to the formation of a functional expression construct where the PolIII promoter of the gRNA gene drives expression of the tag sequences. This usually yields cells with a cytoplasmic fluorescent signal (Fueller et al, 2020). Based on the junction sequences observed across these four major scenarios (Figure 1c), we developed a computational pipeline to analyze insertion site sequencing data and characterize various integration outcomes resulting from PCR tagging (Figure S1a). We applied this pipeline to analyze the outcomes of PCR tagging of one sample case (mNeonGreen tagging of the Gene *PSMD10*) and found that the correct HDR-mediated insertions comprised 59.4% of all detected junctions. NHEJ-mediated junctions at the genomic locus comprise 5.9% whereas NHEJ-mediated concatemeric junctions between two PCR fragments comprise 32.7% of the reads. Fusions between the PCR fragment and non-targeted genomic locations, i.e., the off-target integrations, were sparse (2.0%) (Figure 1f).

To further validate the results, we confirmed that the barcode used in the Tn5-Anchor-Seq had minimal effect on the outcome frequencies. The integration profiles were highly reproducible for the selected target gene across all outcome categories (Figure S1b). The development of this procedure coincided with the transition from ONT R9.4.1/Kit10 to R10/Kit12 chemistry; we thus compared sequencing data generated using both Kits to evaluate the impact of the upgrade. The most notable improvement observed with R10/Kit12 was the longer read lengths, which increased the effective read count available for downstream analysis. With a minimal required length of 290 bp, R9.4.1 chemistry yielded 68.1% of reads above this threshold, while R10 chemistry achieved 92.5% (Figure S1c). Nevertheless, the integration profiles generated by R9 and R10 were highly comparable (Figure S1d).

Given that our customized barcode strategy enables the pooling of a large number of samples for Tn5-Anchor-Seq, we also assessed the rate of index hopping in pooled samples. The observed rates ranged from 0.05% to 6%. Notably, samples with a smaller fraction of total reads exhibited a higher index hopping rate, suggesting that sample proportion may influence the extent of index hopping in pooled sequencing workflows (Figure S1e). Additionally, we observed an agreement between the single-plex sequencing and pooled sequencing with a balanced pool, where samples contribute approximately equal proportion of reads (Figure S1f). This agreement further validates the robustness and reliability of our new methodology.

Taken together we have improved the tagging efficiency using a new selection marker strategy and we developed a quantitative analysis pipeline to assess the outcomes of a tagging experiment in great detail.

### Improvements in tagging efficiency and HDR-mediated integrations

For many applications, it is desirable to obtain populations that are as homogeneous as possible following tagging experiments, avoiding single-clone selection – for example, when a large number of genes need to be tagged. To further enhance the efficiency of precise genome integration via HDR, we next employed our selection strategy and analysis pipeline to explore ways to improve the method further. We aimed to suppress undesired DSB repair outcomes through repression of end-to-end joining of donor dsDNA with the target locus *in vivo* (Figure 2a). For this we tested PCR primers carrying bulky modifications and a stabilized DNA backbone, i.e., biotinylated bases (Btn) and phosphorothioate (PS) coupled bases (Ghanta et al., 2021; Gutierrez-Triana et al., 2018), to generate the PCR cassette. To inhibit NHEJ (Chen et al., 2022; Riesenberg & Maricic, 2018; Robert et al., 2015), we chemically targeted the kinase activity of DNA-PKcs using different inhibitors including KU0060648 (KU), NU7441 (NU), and M3814. Alternatively, we also inhibited ATR (Ataxia telangiectasia mutated and Rad3-related kinase) activity with VE-822 (VE). Concentrations of the drugs were empirically determined (Figure S2a). To suppress MMEJ (Riesenberg et al., 2023), we transiently silenced the Polθ transcript using siRNA (Figure S2b). Finally, we decided to explore the benefits of synchronizing Cas12 expression with the G2 cell-cycle phase to improve HDR efficiency (Gutschner et al., 2016); for this we used an engineered variant, impLbCas12a, fused to the N-terminal region of human Geminin (impLbCas12a-hGEM). impLbCas12a is an improved form of *Lachnospiraceae bacterium* Cas12a (LbCas12a) variant with enhanced nuclease activity and broader range of protospacer-adjacent motif (PAM) sites (Tóth et al., 2020).

**Figure 2.**
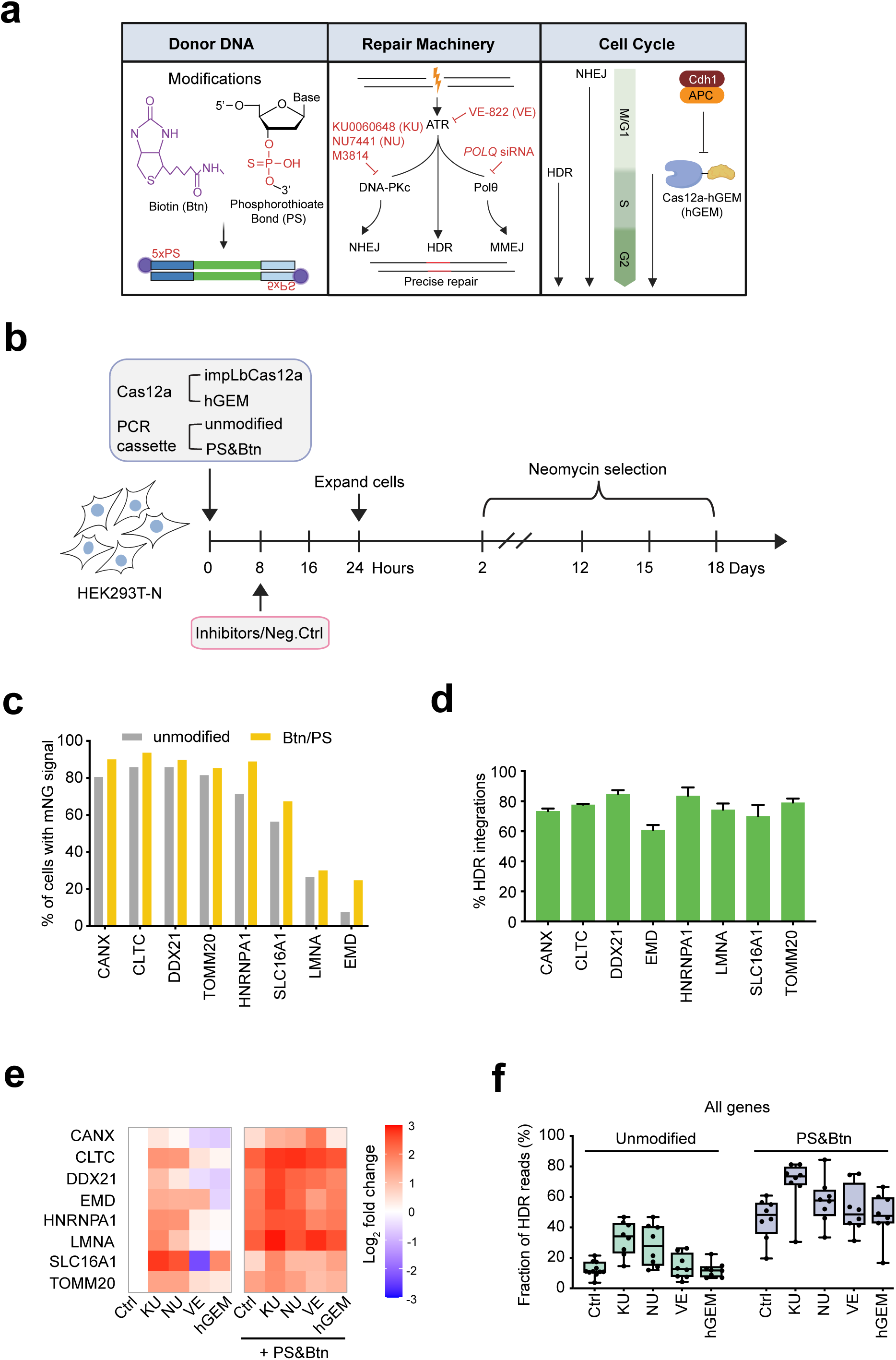
Effects of PCR tagging with various treatments in HDR-mediated integrations. (**a**) Schematic showing the strategies targeting key machineries that are vital to the HDR efficiency of CRISPR/Cas systems. (**b**) Experimental workflow for PCR tagging with various treatments. (**c**) Percentage of cells with mNeonGreen fluorescence at the expected cellular localization, using PCR cassette donors with or without modifications. (**d**) The frequencies of HDR-mediated genomic junctions at various target loci, using PCR cassette donors without modifications. (**e**) Visualization of the improvement of HDR efficiency relative to a tagging experiment with unmodified primers. The heatmap displays log_2_ fold changes in HDR frequency across eight target loci under various treatments, normalized to the condition with unmodified primers and no treatment. (**f**) The average HDR frequency of all target loci in **d**-**e**. Each dot represents the HDR frequency of an individual target, with boxes indicating the interquartile range (25th to 75th percentile), lines denoting the media, and whiskers extending from the minimum to maximum values.

To understand the impact of these conditions over the editing outcome profiles, we selected eight genes and conducted a comparative experiment between several tagging conditions, followed by Tn5-Anchor-seq to obtain data that can quantify their differences in editing fidelity (Figure 2b). We characterized all outcomes using fluorescence microscopy to estimate the tagging efficiency by counting cells with the correct localization of the mNeonGreen tagged protein (mNG positive cells; mNG+), followed by analysis using the ONT sequencing pipeline (Figure 1 and S1). Using the unmodified oligos we observed tagging efficiencies from <10% (EMD) to 80% (Figure 2c). These efficiencies are on average 20-50% higher than previously observed with the constructs where the selection marker is present on the tagging cassette as a separate gene (Fueller et al., 2020). PS- and Btn-modified oligos further improved the efficiencies, with effect sizes in the range of 5% to more than 20%, suggesting that these modifications increase oligo stability, leading to more efficient incorporation during repair. In particular, tagging of the gene with the worst tagging efficiency when using the unmodified oligos (EMD) benefitted the most when using modified oligos.

We also quantified the fraction of mNG+ cells under treatment of NHEJ inhibitors or cell cycle regulated Cas12a, using unmodified or Btn/PS-modified primers. We observed that inhibition of NHEJ with 200nM KU, 1μM NU, or 500nM M3814 increased mNG+ cell proportions across all target genes (Figure S2c,d). However, treatment with 1μM VE-822 reduced mNG+ cell proportions for specific loci, including *DDX21*, *EMD*, *SLC16A1*, and *TOMM20*. Moreover, inhibition of MMEJ with Polθ siRNA for 24h, as well as the use of the cell-cycle-tailored Cas12a-hGEM, did not increase mNG+ cells proportions (Figure S2c,d).

We further analyzed the impacts of these treatments on HDR efficiency using Tn5-Anchor-Seq. We found that with unmodified primers, between 60-85% of reads that involve genomic sequences exhibited the correct insertion by HDR (Figure 2d). Figure 2e visualizes the improvement of HDR efficiency relative to a tagging experiment with unmodified primers, while Figure 2f provides the average HDR efficiency for all genes for each condition. Compared to unmodified PCR cassettes, the frequency of HDR-mediated integration increased by 1.5- to 5.3-fold with PS- and Btn-modified PCR cassettes (Figure 2e, S2e). Treatment with 200nM KU alone largely enhanced HDR efficiency across most genes, although no improvement was observed for *CANX*. Treatment with 1μM NU alone enhanced HDR efficiency for *CLTC*, *EMD*, *HNRNPA1*, *SLC16A1*, and *TOMM20* but not for *CANX*, *DDX21*, and *LMNA*. In contrast, VE-822 and Cas12a-hGEM showed minimal or even negative impacts on HDR efficiency. For all targets, combining PS- and Btn-modified PCR cassettes with NHEJ inhibitors or Cas12a-hGEM resulted in more pronounced improvements in HDR efficiency (Figure 2e, 2f, and S2e). Knockdown of Polθ alone increased HDR efficiency for *EMD* and *SLC16A1,* but not for *LMNA*. However, combining Polθ knockdown with 200nM KU significantly improved HDR efficiency for *LMNA*, while no further improvement was observed for *EMD* and *SLC16A1* (Figure S2g).

Given the high number of individual data points, we did not use technical or biological replicates. Instead, we used the effect of a specific condition on multiple genes as an indicator for its effectiveness. Based on this analysis, we conclude that the most pronounced improvement in HDR efficiency is achieved with the use of modified cassettes, and that efficiency is further enhanced by the addition of NHEJ inhibitors such as KU, NU, and M3814.

### Prevention of unintended indel integrations

In addition to HDR-mediated integration, genome editing can lead to the incorporation of PCR cassettes to the target locus via c-NHEJ pathways. To investigate whether such integration events could be mitigated by the strategies described above (Figure 2a), we performed junction PCR to amplify the insertion junction between the 3’ end of the ORF and the inserted mNeonGreen sequence. This PCR analysis produced two distinct amplicon populations: the shorter band represented junctions formed by HDR-mediated integration, while the longer band corresponded to the expected size of integrations mediated by c-NHEJ (Figure 3a). We quantified the ratio of the HDR band intensity to the total intensity of both bands, defining this as “HDR purity”. At the *HNRNPA1* locus, we observed that both inhibition of c-NHEJ and synchronization of Cas12a increased HDR purity (Figure 3a). We also calculated HDR purity using Tn5-Anchor-Seq data, which showed a strong correlation with the HDR purity obtained from the junction PCR assay across various loci (Figure 3b).

**Figure 3.**
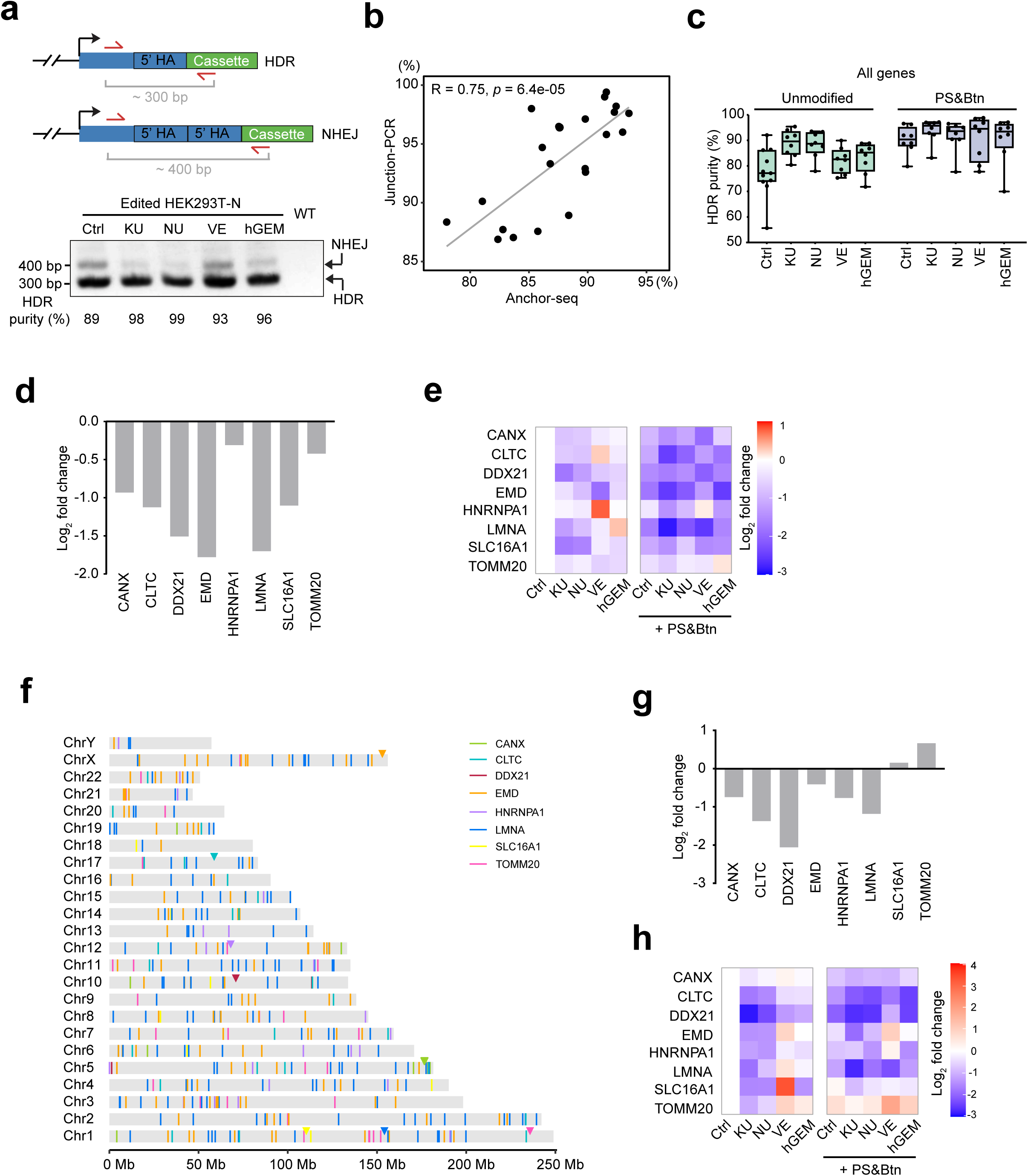
Effects of PCR tagging with various treatments on indel and off-target integrations. (**a**) Schematic showing the insertion outcomes following PCR tagging at the *HNRNPA1* locus and the junction PCR strategy. Red arrows indicate primers binding to the 3’ end of the ORF and the inserted *mNeonGreen* sequence. A representative gel image of junction PCR results and the calculated HDR purity are shown. (**b**) Correlation analysis of HDR purity obtained from junction PCR and Tn5-Anchor-Seq data. Each dot depicts the HDR purity for an individual gene under specific treatment. (**c**) HDR purity of all target loci following PCR tagging with various treatments, as determined by Tn5-Anchor-Seq. Each dot represents the HDR purity of an individual target, with boxes indicating the interquartile range (25th to 75th percentile), lines denoting the media, and whiskers extending from the minimum to maximum values. (**d**) Log_2_ fold changes in indel frequency for eight target loci using PCR cassette donors with PS- and Btn- modifications, relative to unmodified donors. (**e**) Heatmap showing log_2_ fold changes in indel frequency for eight target loci under various treatments, relative to no treatment. (**f**) The distribution of off-target integrations is depicted by colored lines spanning the genome. Triangles of corresponding colors indicate the position of each gene on the respective chromosome. (**g**) Log_2_ fold changes in off-target frequency for eight target loci using PCR cassette donors with PS- and Btn- modifications, relative to unmodified donors. (**h**) Heatmap showing log_2_ fold changes in off-target frequency for eight target loci under various treatments, relative to no treatment.

Analyzing HDR purity across eight genes with Tn5-Anchor-Seq revealed that PS- and Btn-modified PCR cassettes increased HDR purity compared with unmodified cassettes. Treatment with 200 nM KU0060648 or 1 μM NU7441 alone generally enhanced HDR purity for most genes, while 1 μM VE-822 and Cas12a-hGEM showed no obvious effects on HDR purity (Figure 3c, S3a).

We further analyzed the impacts of these treatments on the frequency of NHEJ-mediated indels (in which the mNeonGreen was incorporated with various insertions or deletions within the homology arm and the 3’-ORF region), defined as the ratio of indel read counts to the total genomic integration read counts. Compared to unmodified PCR cassettes, the use of PS and Btn-modified PCR cassettes resulted in a 0.3- to 0.8-fold reduction in indel frequency (Figure 3d). Moreover, combining PS- and Btn-modified PCR cassettes with NHEJ inhibitors or Cas12a-hGEM further reduced indel frequency for all targets except for *TOMM20* (Figure 3e, S3b). Similarly, the combination of PS- and Btn-modified PCR cassettes with Polθ knockdown reduced indel frequency for *EMD* and *SLC16A1*, although no reduction was observed for *LMNA* (Figure S3b).

### Prevention of off-target integrations

In Fueller et al. (2020) we found that off-target integrations resulted from random integration of the PCR cassette independent of Cas12a. Using Tn5-Anchor-seq, we also identified multiple off-target integrations distributed throughout the genome (Figure 3f). To investigate the impacts of the treatments on preventing off-target integrations, we quantified the ratio of the off-target read counts to the total genomic integration read counts, hereafter referred to as the off-target frequency.

The use of PS- and Btn-modified PCR cassettes reduced the off-target frequency by 0.2- to 0.7-fold for most genes but increased the off-target frequency for TOMM20 by 0.6-fold? (Figure 3g). Inhibition of NHEJ with KU, NU, and M3814 generally reduced the off-target integration, while VE-822 increased the off-target integration, especially for *SLC16A1*. Inhibition of MMEJ by knocking down Polθ reduced off-target integration for *EMD* and *SLC16A1* but conversely led to an increase in off-target frequency for *LMNA* (Figure 3h, S3c). These observations are consistent with the idea that inhibition of the DNA repair pathway reduces the frequencies of unwanted events.

### Prevention of concatemeric integration

Consistent with our previous observations (Fueller et al., 2020), a significant proportion of reads containing additional copies of transfected PCR cassettes was detected by Tn5-Anchor-Seq (Figure S3d). These concatemeric reads originate from ligated ends of transfected cassettes, as the free ends of the PCR cassettes are recognized and processed by the c-NHEJ pathway. The most prevalent concatemers are fusions of the PCR cassette, wherein the crRNA gene is ligated to the 5’ end of the mNeonGreen sequence, with the homology arms of the M1 and M2 tagging oligos situated in between (Figure 4a).

**Figure 4.**
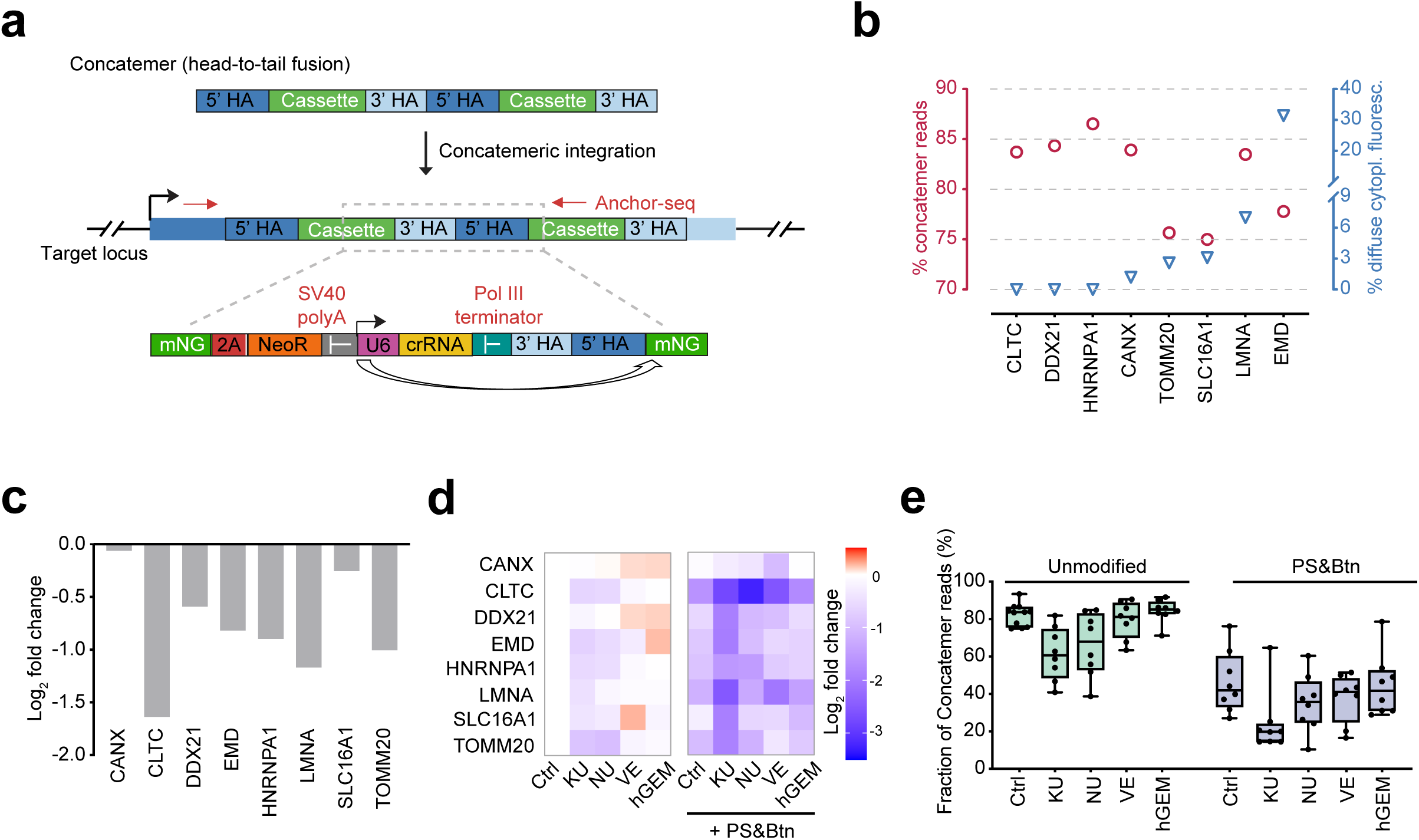
Effects of PCR tagging with various treatments on concatemeric integrations. (**a**) Schematic showing the integration of concatemeric copies of PCR cassettes, highlighting the detailed structure of the tag region. (**b**) Frequency of concatemeric reads and the proportion of cells exhibiting diffuse cytoplasmic fluorescent signals across various loci. (**c**) Log_2_ fold changes in concatemer frequency for eight target loci using PCR cassette donors with PS- and Btn- modifications, relative to unmodified donors. (**d**) Heatmap showing log_2_ fold changes in concatemer frequency for eight target loci under various treatments, relative to no treatment. (**e**) Concatemer frequency of all target loci in **c**-**d**. Each dot represents the concatemer frequency of an individual target, with boxes indicating the interquartile range (25th to 75th percentile), lines denoting the media, and whiskers extending from the minimum to maximum values.

Although an SV40 PolyA terminator sequence was included downstream of the NeoR sequence to ensure proper termination of the gene fusion before the crRNA expression unit, there remains a possibility that the U6 Pol III promoter driving crRNA expression could mediate Pol II-driven expression (Gao et al., 2018; Rumi et al., 2006). This could lead to the expression of mNeonGreen from the subsequent copy in the concatemers (Figure 4a), which partially explains the diffuse cytoplasmic fluorescent signals observed in microscopy in some cells (see examples labeled with white arrows in Figure 2c). We observed that, without any modifications or treatments, 75% - 87% of the reads resulted from concatemers across different loci. This proportion likely overestimates the fraction of concatemer-containing cells, as individual cells may harbor multiple head-to-tail copies. However, no diffuse signals were observed at the *CLTC*, *DDX21*, and *HNRNPA1* loci. Only a few cells (<1%) with diffuse signals were observed from tagging of the *CANX*, *TOMM20*, and *SLC16A1* loci, whereas 7% of cells exhibited diffuse signals at the *LMNA* locus, and 30% of cells showed diffuse signals at the *EMD* locus (Figure 4b).

The PS&Btn modifications on PCR cassettes reduced the frequency of concatemeric reads by 0.5- to 1.7-fold for most genes, while no effect was seen for *CANX* and *SLC16A1* (Figure 4c and Figure S3d). As expected, inhibition of NHEJ with KU, NU, or M3814 decreased the concatemeric integration, while treatment with VE or the use of Cas12a-hGEM increased concatemer frequency at specific loci, such as *CANX*, *DDX21*, and *SLC16A1* (Figure 4d, 4e, and Figure S3e). Inhibition of MMEJ by knocking down Polθ reduced concatemeric integration at the *EMD* and *SLC16A1* loci but not at the *LMNA* locus (Figure S3e).

Our results demonstrate that PS and Btn modifications on PCR cassettes can markedly enhance HDR efficiency while minimizing undesired byproducts, such as NHEJ-mediated indels, off-target integrations, and concatemers. Moreover, combining cassette modifications with pharmacological NHEJ inhibition further improves editing precision, with KU0060648 showing the highest efficacy among the inhibitors tested. Together, these findings establish a robust strategy to improve the accuracy and reliability of PCR-tagging-based genome editing.

### Seamless PCR tagging preserves endogenous 3′-UTR using two PCR cassettes

While the above-described PCR tagging approach (hereafter referred to as *standard PCR tagging*) employed an SV40 PolyA terminator after the tag-sequence to stabilize the transcripts, this editing potentially alters the regulation of the targeted gene. To maintain endogenous post-transcriptional control elements and minimize perturbation of native gene expression, it would be desired to integrate the tag in a seamless manner. This however requires that both ends of the integrated sequence need to be inserted as desired, since aberrant transcripts that would result from the insertion of concatemers could potentially lead to unstable transcripts caused by nonsense-mediated mRNA decay (NMD).

PCR tagging could however also be used for seamless tagging if the PCR fragment with the tag and the marker and the PCR fragment with the gRNA gene are generated separately (see Figure 5a). To test whether this approach works we used the same tagging cassette template plasmid as in the standard PCR tagging workflow and designed new primers to amplify both entities, i.e. the fluorescent protein-2A-neomycin and the gRNA expression construct separately (see Figure 5a). The tag-sequence was designed for precise integration immediately upstream of the stop codon via HDR (and the sequences in the primers M1 and M2*), thereby avoiding introduction of any downstream sequences that could disrupt the endogenous 3′-UTR. In parallel, a separate PCR cassette encoding a guide RNA targeting the desired genomic locus for Cas12a cleavage was amplified using primers M4 and M2*. Both the tag cassette and the guide RNA cassette need to be transfected together with Cas12a expression plasmid.

**Figure 5.**
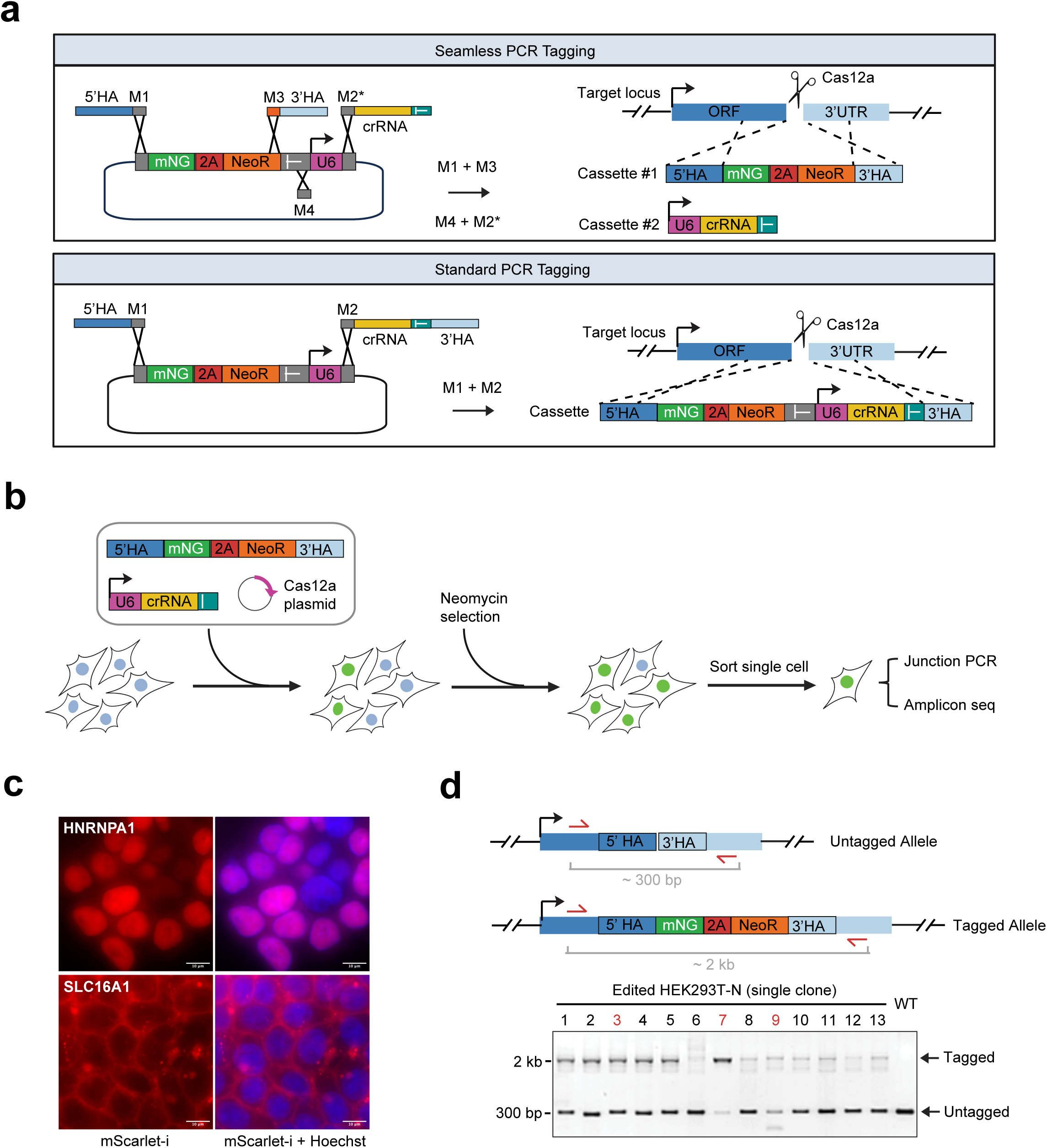
Seamless PCR tagging. (**a**) Schematic of seamless and standard PCR tagging strategies. (**b**) Experimental workflow of seamless PCR tagging and validation: HEK293T-N cells were transfected with two PCR cassettes and Cas12a helper plasmid, followed by neomycin selection and single-cell sorting by flow cytometry. The resulting clonal cell populations were subjected to junction PCR and amplicon sequencing for validation. (**c**) Representative fluorescent microscopy image from HEK293T-N cells showing HNRNPA1 and SLC16A1 tagging. mScarlet-i fluorescence and HOECHST staining are shown. Images acquired using Nikon Ti-E widefield epifluorescence microscope with a 60x objective. Scale bar, 10 µm. (**d**) Schematic showing the insertion outcomes following seamless PCR tagging at the *HNRNPA1* locus and the junction PCR strategy. Red arrows indicate primers binding to the 3’ end of the ORF and the downstream of 3’-UTR. A representative gel image of junction PCR results is shown.

We applied this approach to HEK293T-N cells to insert mScarlet-i-2A-neomycin at the C-terminus of *HNRNPA1* and *SLC16A1*. Fluorescent cells were enriched after 2 weeks of neomycin selection. Images revealed mostly cells with the expected staining (*HNRNPA1*: nucleus; *SLC16A1:* plasma membrane) (Figure 5c). We next isolated clonal cell lines by cell sorting, expanded them and initially verified the insertions by junction PCR using primers flanking the insertion site. This PCR yielded two distinct amplicon sizes: a shorter band corresponding to the untagged allele and a longer band representing the expected size for HDR-mediated integration (Figure 5d). For *HNRNPA1*, 23 single fluorescent clones were isolated, of which 19 (82.6%) displayed both amplicons at their expected sizes. The amplicons of 6 size-correct clones were further analyzed by sequencing, revealing three genotypes in equal proportion: (A) wild-type untagged allele with HDR-tagged allele, (B) untagged allele with small indels plus HDR-tagged allele, and (C) wild-type untagged allele with NHEJ insertion-tagged allele. For *SLC16A1*, 23 (95.8%) out of 24 isolated fluorescent clones possessed both the wild-type and tagged alleles. Further sequencing of 3 size-correct clones showed only Genotype A (Table 2).

**Table 1.** Comparison of standard and seamless PCR tagging methods.

| Feature | Standard PCR tagging | Seamless PCR tagging |
| --- | --- | --- |
| Primer design | One primer pair (M1+M2) | Two primer pairs (M1+M3, M4+M2*) |
| PCR requirement | One PCR reaction | Two PCR reactions |
| Transfection | Cas12a plasmid + one Cassette | Cas12a plasmid + two Cassettes |
| crRNA | On same cassette with donor DNA | Supplied separately |
| 5'-UTR | Endogenous 5'-UTR | Endogenous 5'-UTR |
| 3'-UTR | Exogenous 3'-UTR | Endogenous 3'-UTR |

**Table 2.** Characterization of individual fluorescent clones following seamless PCR tagging.

| Gene | Single fluorescent clones | Size-correct Junction PCR | Amplicon Sequencing | Genotype<sup>*</sup> |  |  |
| --- | --- | --- | --- | --- | --- | --- |
|  |  |  |  | A | B | C |
| **HNRNPA1** | 23 | 19 | 6 | 2 | 2 | 2 |
| **SLC16A1** | 24 | 23 | 3 | 3 | 0 | 0 |

\*: Genotype A: Wildtype untagged allele + HDR-tagged allele (Desired edit); Genotype B: Untagged allele with small indels + HDR-tagged allele; Genotype C: Wildtype untagged allele + NHEJ-tagged allele. Please note that HEK293 cells are partially polyploid (according to ATCC) and the exact copy number of individual chromosomes in our cells is not known.

This demonstrates that PCR tagging can be used for seamless tagging. For convenient primer design, we have extended the functionality of the PCR-tagging.com website. This resource, originally developed for standard PCR tagging primer design, has now been expanded to also support seamless tagging.

## Discussion

Here, we advance mammalian PCR tagging (Fueller et al., 2020) on two fronts: (i) a practical set of design and workflow changes that elevate the fraction of precise, HDR-mediated insertions while suppressing common by-products; and (ii) a sequencing and analysis framework (Tn5-Anchor-Seq coupled to a reproducible pipeline) that resolves the full spectrum of on-target and off-target outcomes generated by CRISPR/Cas12a–assisted PCR tagging. Together, these developments turn our previous approach into a scalable, higher-fidelity strategy for routine gene tagging, and they provide a quantitative lens for mechanistic optimization.

A central finding is that chemically stabilizing the PCR donor ends with phosphorothioate bonds and terminal biotin, combined with a selection marker fused in-frame via a 2A peptide to enrich for expressed cells, shifts repair toward HDR and away from end-joining outcomes. Across diverse loci, these cassette designs consistently increased the proportion of correctly localized fluorescent signals and the fraction of HDR junctions captured by sequencing. Pharmacological inhibition of DNA-PKcs (KU0060648, NU7441, M3814) further amplified this effect and, when paired with modified donors, produced the most pronounced gains in HDR and reductions in indels, concatemers, and off-target integrations. In contrast, ATR inhibition provided little benefit and sometimes harmed performance, underscoring that generic “HDR-promoting” interventions can have pathway- and locus-specific liabilities. Likewise, transient suppression of Polθ (MMEJ) produced modest or gene-contingent improvements in our context, in line with emerging evidence that the relative contributions of c-NHEJ, MMEJ, and HDR vary with sequence architecture, chromatin environment, and cell state. Cell-cycle tailoring of Cas12a activity (hGEM-Cas12a fusion) yielded at most incremental gains compared with donor end-protection plus DNA-PKcs inhibition, suggesting that repairing the substrate (the donor ends) and disfavoring the dominant end-joining pathway are the highest-leverage levers for cassette-based knock-ins.

These mechanistic trends are mirrored in the classes of by-products we quantified. Concatemeric insertions—often invisible to short-amplicon assays—were frequent without intervention and could explain diffuse cytoplasmic fluorescence in a locus-dependent manner. Modified donors and DNA-PKcs inhibition reduced concatemer frequency, consistent with protection of donor ends from ligation and a general dampening of c-NHEJ activity. Off-target genomic integrations, which arise independently of the programmed cut, also fell under these conditions. Importantly, our sequencing and classification approach distinguishes HDR junctions at the intended locus from (i) imperfect but in-frame NHEJ fusions that can still produce localized fluorescence; (ii) concatemer junctions between donor copies; and (iii) off-target insertions elsewhere in the genome. This resolves a known ambiguity of selection-based readouts: a cell can be drug-resistant and fluorescent for reasons other than a perfect on-target HDR event. The improved 2A-linked selection enriches for correctly tagged fusions but does not, by itself, guarantee perfect structure; hence the value of orthogonal molecular auditing.

Methodologically, Tn5-Anchor-Seq brings several practical advantages. Library construction from genomic DNA is straightforward and scalable: Tn5 tagmentation, biotin-assisted capture of anchor-proximal fragments, and nested PCR collectively enrich insert–genome junctions with high specificity. The long reads achievable on ONT provide contiguous visibility across junction complexity (microindels, duplications within homology arms, donor–donor fusions), while sample barcoding enables efficient pooling without compromising concordance. The accompanying Snakemake pipeline automates basecalling-to-classification with transparent, reproducible rules, which should facilitate adoption and cross-study comparisons.

At the same time, several limitations temper interpretation. First, because the assay targets junctions rather than whole alleles, it does not yield an absolute copy number of donor concatemers; very long arrays and integrations that delete the anchor region can be undercounted or missed. Second, ONT error profiles (particularly indels at homopolymers) can blur the precise microstructure of individual repair events; we mitigate this by focusing on categorical outcomes and by validating trends across loci, but single-event reconstructions should be interpreted cautiously. Third, our optimization screen prioritized breadth (multiple loci and conditions) over depth (replicates), using gene-to-gene consistency to judge effect sizes. Although the directional effects are robust, formal replication in additional cell types—especially primary and therapeutically relevant cells—will refine dose windows for inhibitors and generalize the parameter space. Finally, selection based on a 2A-linked marker improves the fraction of functional fusions but could, in principle, bias against edits that affect translation or protein stability; users should choose markers with the biology of the target in mind.

We introduce a **seamless PCR tagging** variant that structurally preserves endogenous 3′-UTRs by supplying the crRNA cassette separately from the tag donor. This design eliminates exogenous poly(A) signals from the edited locus and better maintains native post-transcriptional regulation. In clonal analyses, the approach produced a high proportion of size - consistent HDR alleles with expected localization, though - as for any DSB-based editing - minor fractions of NHEJ-containing or mixed genotypes persisted. The trade-offs are practical: two PCRs and co-delivery of an additional cassette modestly increase workflow complexity and, theoretically, add routes for donor–donor ligation. Nonetheless, for genes where 3′-UTR integrity matters, the benefits outweigh these costs, and our results show that seamless tagging can be executed with the same donor-protection and pathway-modulation principles to maintain high fidelity.

Looking forward, several refinements could further tighten precision and interpretability. On the sequencing side, unique molecular identifiers (UMIs) at the tagmentation step, duplex-consensus polishing, and complementary short-read capture of the opposite junction would improve quantitation and error correction, and could enable approximate concatemer copy number estimation. On the editing side, combining donor end-protection with transient, low-dose DNA-PKcs inhibition appears broadly useful; future work should map minimal effective exposures across cell types to balance efficacy and cytotoxicity. Incorporating insulators or transcriptional terminators flanking Pol III promoters may further suppress spurious expression from concatemers. Finally, the Tn5-Anchor-Seq logic is nuclease-agnostic and payload-agnostic; it should generalize to Cas9, nickase-based strategies, transposase or integrase systems, and larger functional cassettes (e.g., degrons, barcodes), providing a common measurement scaffold for systematic optimization.

In sum, by coupling rational cassette engineering and repair-pathway modulation with a long-read junction-profiling toolkit, we deliver a practical, cloning-free route to higher-fidelity mammalian gene tagging and a means to audit what actually happened at the genome level. We anticipate that these advances will lower the barrier to large-scale tagging efforts, enable cleaner pooled or population-level experiments without extensive clonal isolation, and improve the reliability of downstream functional genomics in mammalian systems.

## Methods

### Cell culture

The neomycin sensitive HEK293T (HEK293T-N) cells (gift from Jonathan Schmid-Burgk, University of Bonn, Germany) were cultured in DMEM high glucose (Life Technologies) supplemented with 10% (vol/vol) FBS (Gibco). Cells were cultured at 37°C with 5% CO_2_ and regularly tested for mycoplasma contamination.

### Generation of Plasmids and PCR cassettes

The pMaCTag-2A-N07 template plasmid for PCR cassette amplification was constructed using plasmid #9697 (this plasmid was kindly provided by Jonathan Schmid-Burgk (Zhang et al., 2021: Zhang F, Schmid-Burgk JL, Hornung V. Sequencing-based proteomics. US-Patent US20210147831/US12281301.)) and plasmid pMaCTag-Z07 (Addgene #124786). Specifically, the mNeonGreen-T2A-NeoR-SV40polyA sequence was PCR amplified from plasmid #9697, digested with BamHI and XhoI, and subsequently subcloned into plasmid pMaCTag-Z07, which was pre-digested with the same enzymes.

The pMaCTag-2A-N11 template plasmid was constructed as follows: the mScarlet-i fragment was excised from pMaCTag-11 (Addgene #119990) with BamHI and BsrGI, then subcloned into plasmid pMaCTag-2A-N07 that was pre-digested in the same way.

The cell-cycle-tailored Cas12a helper plasmid (pcDNA3.1-impLbCas12a-hGem) was constructed as follows: the hGem1-110 fragment was amplified from RPE-1 cDNA, cut with BamHI and EcoRI, then subcloned into plasmid pVE13300 (Addgene #137715) that was pre-digested in the same way. All restriction enzymes were purchased from NEB.

Tagging oligos (M1, M2, M2*, M3, and M4) used for generating PCR cassettes were designed using the online tool (www.pcr-tagging.com) and obtained from Sigma-Aldrich with RP1 cartridge purification. The modified oligos consist of the same sequences as the unmodified ones but include phosphorothioate bonds in the first five nucleotides and have a biotin modification at the 5’ end. The PCR cassettes were amplified from either pMaCTag-2A-N07 (mNeonGreen) or pMaCTag-2A-N11 (mScarlet-i) template plasmid based on our previously published protocol (Fueller et al., 2020). All the primers and oligos used for cloning, validation and PCR cassettes are listed in Table S1.

### Transfection

The transfection of HEK293T-N cells was performed in a 24-well format with Lipofectamine 3000 (Invitrogen) according to the manufacturer’s protocol. Briefly, cells at ∼70% confluency in one well of a 24-well plate were transfected with 100 ng of Cas12a helper plasmid (either impLbCas12a (Addgene #137715) or impLbCas12a-hGem) and 900 ng PCR cassette. One day after transfection, cells were expanded to a 6-well plate. If NHEJ inhibitors were applied, cell culture media containing small molecules (KU0060648, NU7441, VE-822, M3814) were added after 8 hours of transfection and cells were incubated for 16 hours. Two days after transfection, cells were selected with 500 µg/ml neomycin (Sigma-Aldrich).

If applicable, 30pmol of siRNA765 duplexes (DsiRNA hs.Ri.Polθ.13.8, Integrated DNA Technologies) were transfected together with Lipofectamine 3000 for knockdown Polθ. a non-targeting siRNA was used as a negative control. Cells were incubated for 8-24 hours before downstream experiments.

### Cell viability measurement

Cell viability was measured using the CellTiter-Blue Cell Viability Assay (Promega TB317) according to the manufacturer’s instructions. HEK293T-N cells were seeded in a 96-well plate at a density of 10^4^ cells per well. Cells were treated with varying concentrations of NHEJ inhibitors (KU0060648, NU7441, VE-822, M3814) for different time points. After 24 h of treatment, 20 µl of CellTiter-Blue reagent was added to each well containing 100 µl of culture medium. Plates were incubated at 37°C for 2h to allow resazurin reduction, and fluorescence was measured at 560 nm excitation and 590 nm emission with Tecan plate reader (Tecan Spark Cyto). Background fluorescence from medium-only wells was subtracted, and viability was expressed relative to untreated control.

### Fluorescence microscopy

Cells were seeded in 8-well chamber slides and incubated for 4 hours before imaging. Cells were stained with Hoechst 33342 (4 µg/ml in PBS, Thermo Fisher Scientific) for 10 min. Images were acquired on a Nikon Ti-E widefield epifluorescence microscope with a 60x ApoTIRF oil-immersion objective (1.49-NA(numerical aperture); Nikon), an LED light engine (SpectraX, Lumencor), a 2048 x 2048 pixel (6.5 µm) sCMOS camera (Flash4, Hamamatsu) and an autofocus system (Perfect Focus System, Nikon) with either bright field or 469/35-nm excitation filter (Semrock) and 525/50-nm emission filter (Chroma). Z-stacks of 11 planes with 0.5-µm spacing were recorded with 100-ms exposure time for all cells, maximum intensity z-projections are shown. Subcellular localizations were identified and scored visually. For cell counting, random fields of view were inspected in the Hoechst channel, and all nuclei present in the entire field were counted.

### Fluorescence-activated cell sorting

Tagged cells were harvested, washed with PBS, and resuspended at a density of 10^6^ cells/ml in PBS supplemented with 1% BSA (BioShop, ALB001). Single-cell sorting was performed on a Aria Fusion or Symphony S6 flow cytometer (BD Biosciences) with nozzle size of 100 µm. Forward scatter and side scatter parameters were used to discriminate intact cells from debris. mScarlet-i intensity was measured in the G575 or Y586 channels and used as the sorting parameter. Non-viable DAPI positive cells were excluded using the V450/V431 channel. mScarlet-i-positive cells were sorted directly into 96-well plates containing DMEM high glucose media supplemented with 10% (vol/vol) FBS and 1% penicillin-streptomycin.

### Genomic DNA isolation

Genomic DNA (gDNA) was extracted from samples using Monarch Genomic DNA purification kit (NEB, T3010) according to the manufacturer’s protocol. Briefly, cell pellets containing 5 x 10^5^ cells were resuspended in 100 µl of cold PBS and incubated with 100 µl of Cell Lysis Buffer containing Proteinase K and RNase A at 56 °C for 5 minutes. The lysates were then mixed with 400 µl Binding Buffer and loaded onto gDNA Purification Columns to bind the gDNA. The flow-through was discarded, and the columns were washed twice with 500 µl of Wash Buffer. Finally, the gDNA was eluted in 60 µl of preheated (60 °C) Elution Buffer. After elution, the gDNA concentration was measured with NanoDrop, and its integrity and size were assessed by gel electrophoresis on 1% agarose/TAE.

### Junction PCR and amplicon sequencing

To check for the integration of the PCR cassettes, PCR amplification of the insertion junction from the wide-type and tagged cells was performed with Velocity polymerase (Bioline) or Platinum Direct PCR Universal Master Mix (Thermo Scientific). For standard PCR tagging, the primers specific to mNeonGreen sequence and gene-specific primers annealing outside of the 5’-homology arms of the target gene were used. The PCR products were analyzed by gel electrophoresis on 1% agarose/TAE, where the upper band corresponds to junctions generated by integration via c-NHEJ and the lower via HR. For seamless PCR tagging, the primers bind to the 3’-end of the ORF and the downstream of 3’-UTR. The PCR products were analyzed by gel electrophoresis on 1% agarose/TBE. For sequencing validation, the PCR products were purified with Agencourt AMPure XP beads (Beckman) according to the manufacturer’s instructions. Premium PCR Sequencing was performed by Plasmidsaurus using Oxford Nanopore Technology. The primers used for junction PCR are listed in Table S1.

### Tn5-Anchor-Seq library preparation and ONT sequencing

Sequencing libraries for Tn5-Anchor-Seq were prepared following our previously published Anchor-Seq protocol with some modifications(Fueller et al., 2020). In brief, 300 ng of Tn5 (E54K, L372P, 50 ng/µl) transposase (Hennig et al., 2018) was preloaded with 1.5 µM annealed adapters (P5-UMI-xi5001…5096-ME.fw, Tn5hY-Rd2-Wat-SC3) in 50 mM Tris-HCl (pH 7.5) at 23 °C for 1h. The preloaded transposase was mixed with 400 ng of gDNA, 10 mM Tris-HCl (pH 7.5), 10mM MgCl2, and 25%[v/v] dimethylformamide, and incubated at 55 °C for 10min to tagment the gDNA. Tagmented gDNA was purified using NGSPrep beads (Steinbrenner, MDKT00010005) according to the manufacturer’s instructions. The purified tagmented gDNA was used as a template for a first PCR reaction with a biotinylated cassette-specific primer and Tn5 adapter-specific primer, using LongAmp Hot Start Taq2 polymerase (NEB, M0533). The reaction was run for 15 cycles with 63 °C annealing and 70s elongation. Biotinylated amplicons were initially purified using NGSPrep beads, followed by enrichment with Dynabeads MyOne Streptavidin C1 beads (Invitrogen, #65001). These beads were used as the input of a second PCR, which used nested cassette- and Tn5 adapter-specific primers with LongAmp Hot Start Taq2 polymerase. The following program was used: 3 min at 95 °C; 5 cycles of 30s at 95 °C, 60 s at 60 °C, and 70 s at 65 °C; 30 cycles of 15s at 95 °C, 15 s at 62 °C, and 70 s at 65 °C; 5 min at 65 °C; and 4 °C hold. PCR products longer than 500bp were size-selected using NGSPrep beads and subsequently ligated with ONT sequencing adapter to generate the final library. The SQK-PBK004 was used for R9.4.1 flow cells, while the SQK-LSK114 kit was used for R10.4.1 flow cells. Libraries were quantified using Qubit, loaded onto the flow cell according to the manual, and run on the MinION device (ONT). The primers used for Tn5-Anchor-Seq are listed in Table S1.

### Integration outcome analysis

The raw signal data were generated in either Fast5 (R9.4.1) or Pod5 (R10.4.1) formats and basecalled to FASTQ files using Dorado v0.8.1 (ONT). Quality assessment of the reads was performed with NanoPlot v1.41.6. Reads were then processed using custom scripts in Julia v1.8.3 with the BioSequences v2.0.1 package for demultiplexing by barcodes and removal of Tn5 adapter and constant sequences. The demultiplexed and trimmed reads were first aligned in ONT mode using bwa-mem v0.7.17-r1188 against a custom reference genome (Li, 2013). This custom reference consisted of both wild-type and PCR cassette sequences for all target genes, with the wild-type sequence defined as up to 10,000 bases (depending on the gene length) upstream of the 5’-homology regions for each target gene. The aligned reads were filtered and annotated as HDR, Indel, or concatemer reads using in-house Python scripts. Unaligned reads and reads with low mapping quality were further aligned against the hg38 human genome to identify potential off-target integrations. Identified integration outcomes were evaluated using IGV v2.16.1 and further quantified by custom R scripts. All custom scripts were integrated into a Snakemake workflow, allowing automated execution of the analysis pipeline.

## Data availability

The data generated in this study are available in this article and its supplementary data files. The computational pipeline for characterizing and quantifying genomic integration outcomes is available on GitHub at https://github.com/dlou2022/PCRtaggingProfiler.

## Acknowledgments

We thank Jonathan Schmid-Burgk for providing cell lines and plasmids and for helpful discussions throughout this work. We are grateful to Simon Anders for support with the design and analysis of the sequencing experiments. We also thank the members of our laboratories for critical reading of the manuscript and for discussions.

This work was supported by the Baden-Württemberg Stiftung through the project MET-ID53, funded within the program *Methoden in den Lebenswissenschaften*

