## supplemental figures for "Improved cloning-free one-step CRISPR-Cas12a-assisted tagging of mammalian genes using PCR generated reagents (PCR tagging)" (from another version of this preprint: version 1, 2025-12-02)

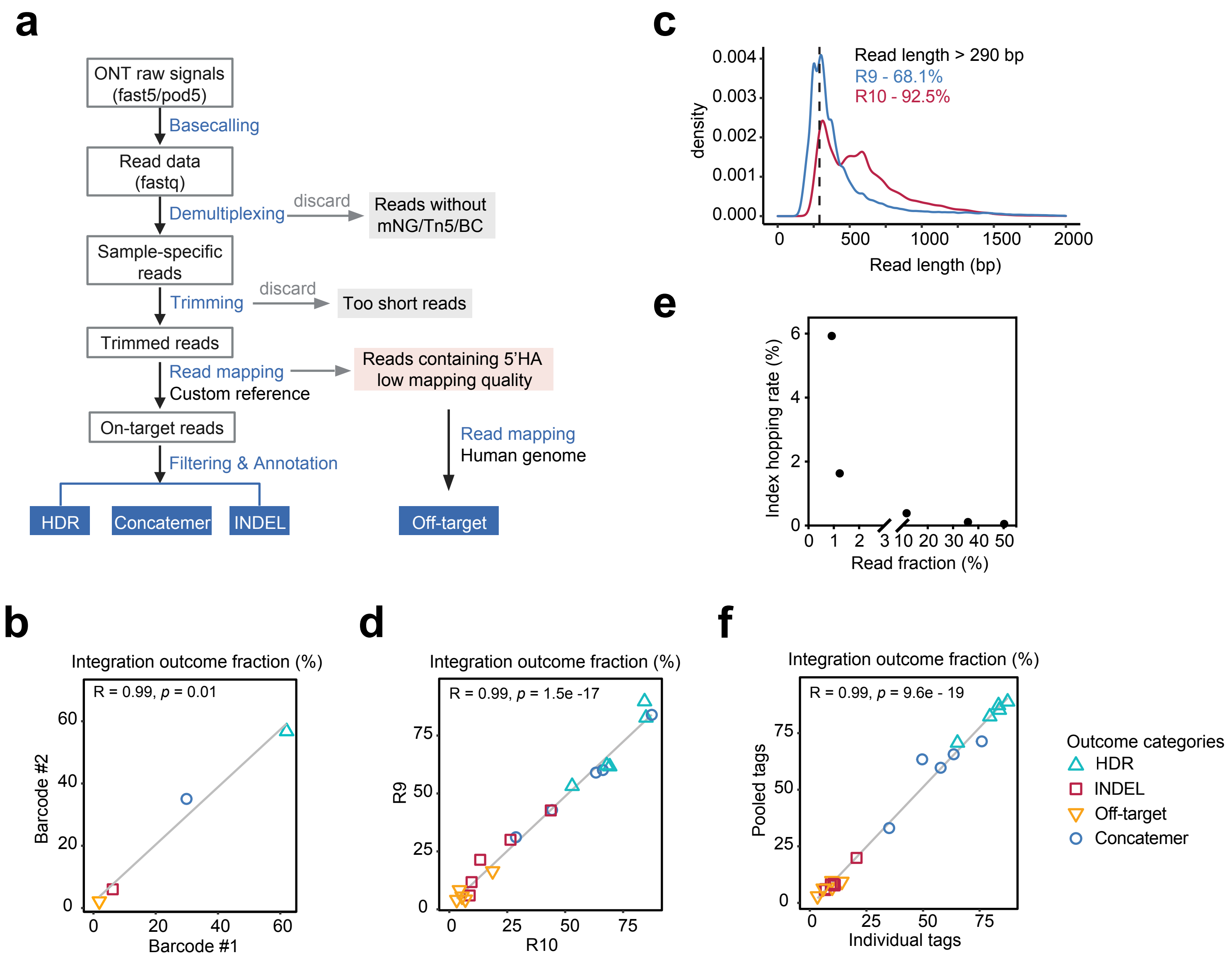

a
ONT raw signals (fast5/pod5)
Basecalling
Read data (fastq)
Demultiplexing
discard
Reads without mNG/Tn5/BC
Sample-specific reads
Trimming
discard
Too short reads
Trimmed reads
Read mapping
Custom reference
Reads containing 5'HA low mapping quality
On-target reads
Read mapping
Human genome
Filtering & Annotation
HDR
Concatemer
INDEL
Off-target
b
Integration outcome fraction (%)
R = 0.99, p = 0.01
Barcode #2
Barcode #1
c
Read length > 290 bp
R9 - 68.1%
R10 - 92.5%
density
Read length (bp)
d
Integration outcome fraction (%)
R = 0.99, p = 1.5e -17
R9
R10
e
Index hopping rate (%)
Read fraction (%)
f
Integration outcome fraction (%)
R = 0.99, p = 9.6e - 19
Pooled tags
Individual tags
Outcome categories
HDR
INDEL
Off-target
Concatemer

**Figure S1**. **Computational classification pipeline to characterize genomic integration outcomes resulting from PCR-tagging, related to Figure 1**

(**a**) Detailed data processing pipeline of Tn5-Anchor-Seq. (**b**) The frequencies of integration outcomes at PSMD10 locus compared between different barcodes. Each point depicts the fraction of individual outcome. (**c**) Read length distribution of Tn5-Anchor-Seq data generated by R9.4.1/Kit10 (R9) and R10/Kit12 (R10) chemistries. (**d**) The frequencies of integration outcomes at various target loci compared between R9 and R10 chemistries. Each point depicts the fraction of individual outcome. (**e**) Index hopping rates and read fractions of pooled samples. (**f**) The frequencies of integration outcomes at various target loci compared between single-plex and pooled sequencing. Each point depicts the fraction of individual outcome.

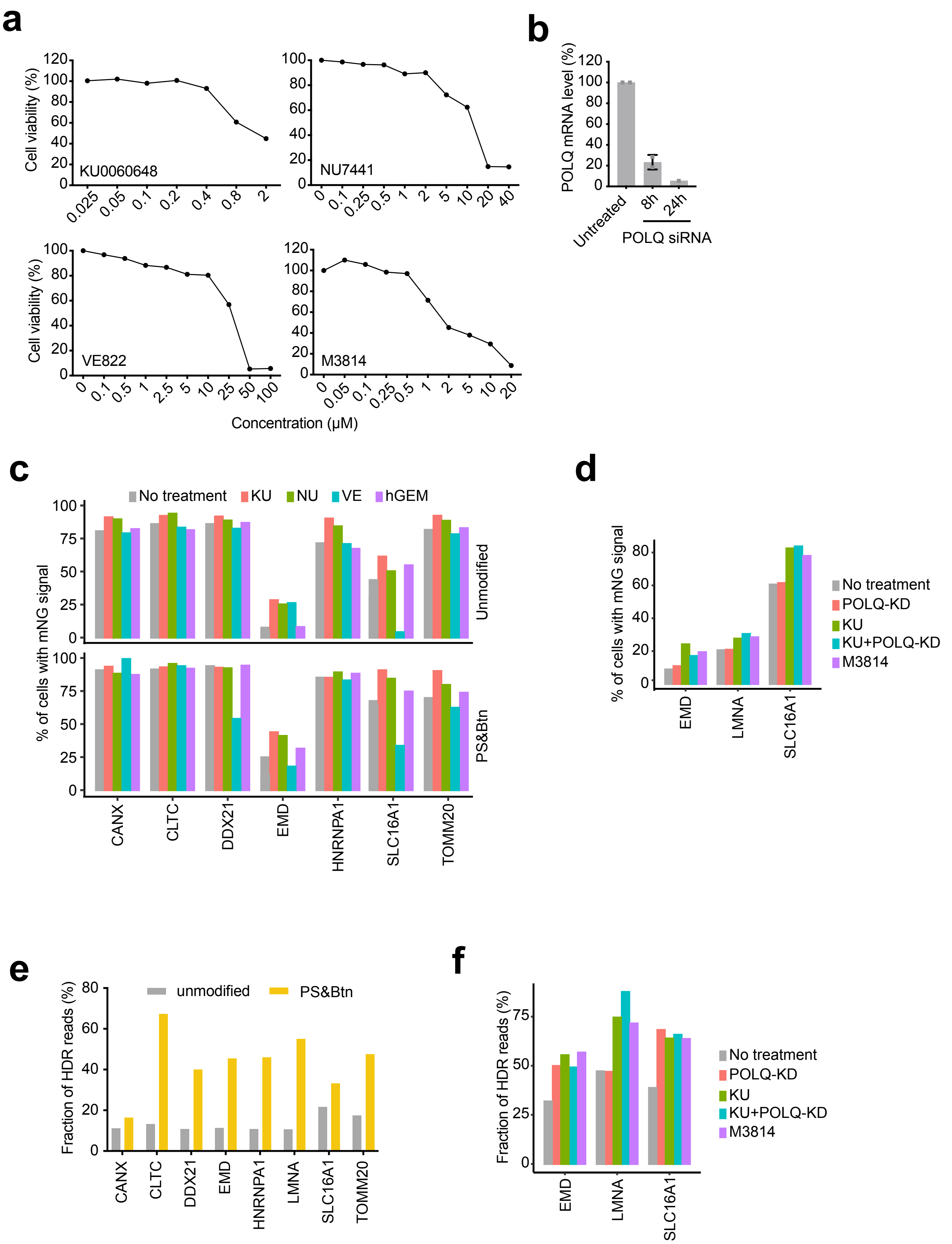
a
Cell viability (%)
KU0060648
NU7441
VE822
M3814
Concentration (μM)
b
POLQ mRNA level (%)
Untreated
8h
24h
POLQ siRNA
c
No treatment
KU
NU
VE
hGEM
% of cells with mNG signal
Unmodified
PS&Btn
CANX
CLTC
DDX21
EMD
HNRNPA1
SLC16A1
TOMM20
d
% of cells with mNG signal
No treatment
POLQ-KD
KU
KU+POLQ-KD
M3814
EMD
LMNA
SLC16A1
e
Fraction of HDR reads (%)
unmodified
PS&Btn
CANX
CLTC
DDX21
EMD
HNRNPA1
LMNA
SLC16A1
TOMM20
f
Fraction of HDR reads (%)
No treatment
POLQ-KD
KU
KU+POLQ-KD
M3814
EMD
LMNA
SLC16A1

**Figure S2**. **Cell viability of HEK293T-N cells treated with small molecule inhibitors (KU0080648, NU7441, VE-822, and M3814) and tagging efficiency after PCR tagging with the treatments. (a**) The HEK293T-N cells were treated with various concentrations of inhibitors for 24 hours, and cell viability was determined in relative to vehicle controls (N=3). (**b**) Time course of Polθ mRNA knockdown in HEK293T-N cells with siRNA765. Polθ expression was normalized with *GAPDH* expression and is shown relative to untreated cells. (**c**) Percentage of cells with mNeonGreen fluorescence at the expected locus following PCR tagging with treatments involving KU0060648, NU7441, VE-822, or Cas12a-hGEM. All PCR cassettes used in this experiment were modified with PS and Btn. (**d**) Percentage of cells with mNeonGreen fluorescence at the expected locus following PCR tagging with Polθ knockdown or treatments with KU0060648 or M3814. All PCR cassettes used in this experiment were modified with PS and Btn. (**e**) HDR frequency following PCR tagging, using PCR cassette donors with or without PS- and Btn- modifications. (**f**) HDR frequency following PCR tagging with Polθ knockdown or treatments with KU0060648 or M3814. All PCR cassettes used in this experiment were modified with PS and Btn.

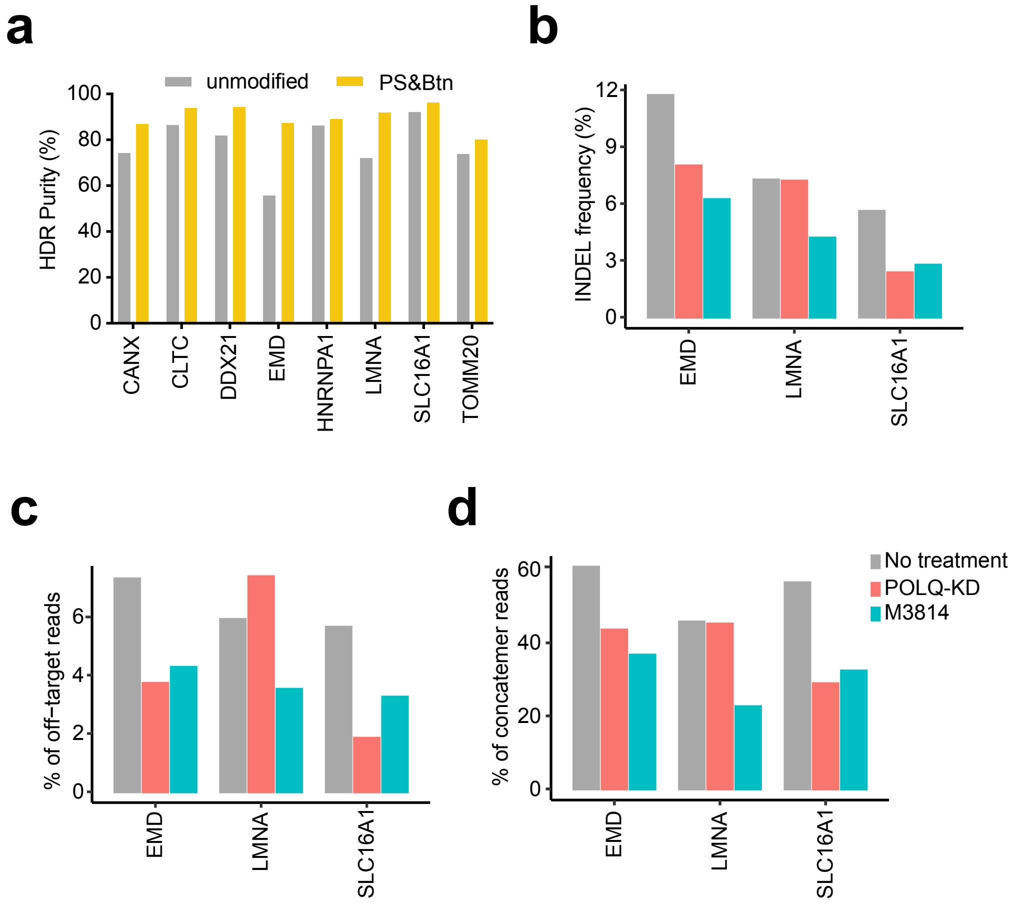

a
unmodified
PS&Btn
HDR Purity (%)
100
80
60
40
20
0
CANX
CLTC
DDX21
EMD
HNRNPA1
LMNA
SLC16A1
TOMM20
b
INDEL frequency (%)
12
9
6
3
0
EMD
LMNA
SLC16A1
c
% of off-target reads
6
4
2
0
EMD
LMNA
SLC16A1
d
% of concatemer reads
60
40
20
0
EMD
LMNA
SLC16A1
No treatment
POLQ-KD
M3814

**Figure S3**. **Indel, off-target, and concatemeric integrations following PCR tagging under various treatments. (a**) HDR purity following PCR tagging, using PCR cassette donors with or without PS- and Btn- modifications. (**b**) Indel frequency following PCR tagging with *POLQ* knockdown or treatments with M3814. All PCR cassettes used in this experiment were modified with PS and Btn. (**c**) Off-target frequency following PCR tagging with Polθ knockdown or treatments with M3814. All PCR cassettes used in this experiment were modified with PS and Btn. (**d**) Concatemer frequency following PCR tagging with Polθ knockdown or treatments with M3814. All PCR cassettes used in this experiment were modified with PS and Btn.
